# Simultaneous single cell measurements of intranuclear proteins and gene expression

**DOI:** 10.1101/2021.01.18.427139

**Authors:** Hattie Chung, Christopher N. Parkhurst, Emma M. Magee, Devan Phillips, Ehsan Habibi, Fei Chen, Bertrand Yeung, Julia Waldman, David Artis, Aviv Regev

## Abstract

Identifying gene regulatory targets of nuclear proteins in tissues remains a challenge. Here we describe <u>in</u>tranuclear <u>C</u>ellular <u>I</u>ndexing of <u>T</u>ranscriptomes and <u>E</u>pitopes (inCITE-seq), a scalable method for measuring multiplexed intranuclear protein levels and the transcriptome in parallel in thousands of cells, enabling joint analysis of TF levels and gene expression *in vivo*. We apply inCITE-seq to characterize cell state-related changes upon pharmacological induction of neuronal activity in the mouse brain. Modeling gene expression as a linear combination of quantitative protein levels revealed the genome-wide effect of each TF and recovered known targets. Cell type-specific genes associated with each TF were co-expressed as distinct modules that each corresponded to positive or negative TF levels, showing that our approach can disentangle relative contributions of TFs to gene expression and add interpretability to gene networks. InCITE-seq can illuminate how combinations of nuclear proteins shape gene expression in native tissue contexts, with direct applications to solid or frozen tissues and clinical specimens.

## Main Text

Single nucleus RNA sequencing is an essential tool for profiling the heterogeneity of solid tissues, particularly for clinical studies of tissues that are challenging to dissociate or frozen, or for studies that require preserving cellular activity states by avoiding non-specific activation induced by enzymatic cellular dissociation^1–7^. As the nucleus is also a key site of gene regulation by proteins such as transcription factors (TFs), simultaneously measuring proteins and the transcriptome inside individual nuclei would enable integrating rich phenotypic and genomic information in tissues. Methods using DNA-conjugated antibodies to measure surface proteins and RNA at single cell resolution, such as CITE-seq^8^, have been widely applied to circulating immune cells^9, 10^, and recently to cytoplasmic protein targets^11, 12^, but they are less suited for non-immune cells and tissues where dissociation disrupts the integrity of cellular membranes. A single nucleus-based method for measuring quantitative protein levels and the transcriptome is an open challenge, particularly as DNA-conjugated antibodies are “sticky” inside the nucleus due to ubiquitous non-specific binding^13^.

Simultaneously measuring nuclear proteins and RNA in single cells would enable relating nuclear levels of proteins to newly transcribed RNA to reveal genes and pathways involved in cell state changes^2, 14–18^, and how gene networks regulated by TFs vary across context and in disease^19^. Aberrations in nuclear levels of specific TFs can be hallmarks of disease and even predict patient outcomes^20–24^. Furthermore, nuclear localization of TFs can indicate a change in cellular activity states such as in the case of TFs that shuttle in and out of the nucleus in response to environmental signals^25–30^. For example, the activity-regulated transcription factor complexes NF-κB and AP-1, and their components p65 and c-Fos, transiently localize to the nucleus in response to different signals^31–34^, where they activate the expression of diverse pathways related to inflammation, oncogenesis, apoptosis, cell proliferation, and synaptic remodeling^35–43^.

Current studies that monitor nuclear TF levels and gene expression typically rely on live cell imaging in culture, *in situ* measurement of proteins and RNA in tissue by staining and hybridization, or cell sorting and profiling based on fluorescent reporters. Such studies have shown that due to asynchrony in responses and dynamic shuttling, nuclear localization can vary between individual cells stimulated together^44, 45^. However, these methods relying on reporters or a handful of probes are limited in their ability to relate changes in protein localization to their genome-wide impact on transcription.

Here we report intranuclear CITE-seq (“inCITE-seq”), a method that enables multiplexed and quantitative intranuclear protein measurements using DNA-conjugated antibodies coupled with RNA sequencing on a droplet-based sequencing platform (**Fig. 1a**). To allow antibody diffusion across the nuclear membrane, nuclei are lightly fixed with formaldehyde and permeabilized, blocked under optimized conditions to minimize non-specific binding of DNA-conjugated antibodies inside the nucleus, and combined with nucleus hashing antibodies^46^ to allow multiplexing (**Methods**). We then load antibody-stained nuclei directly for droplet-based snRNA-seq for simultaneous capture of antibody DNA tags and the transcriptome. We demonstrate the utility of inCITE-seq for profiling response to environmental stimuli in cells and tissues.

**Figure 1.**
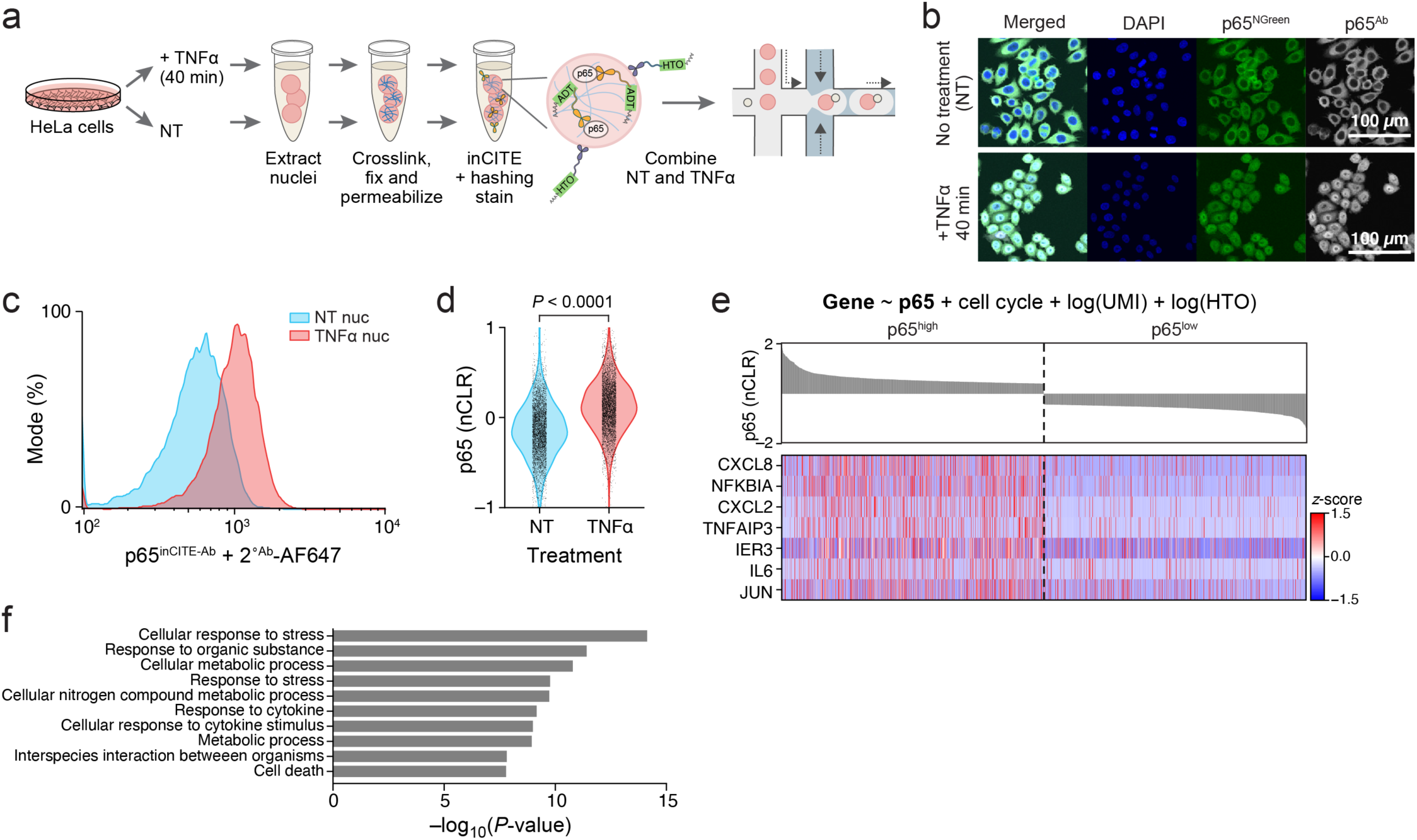
InCITE-seq for simultaneous measurement of intranuclear protein and RNA levels at single nucleus resolution. **a.** InCITE-seq method overview in HeLa cells for droplet-based nucleus profiling. **b.** TNF*α* induced nuclear translocation of p65. Fluorescent imaging of HeLa cells expressing a p65- mNeonGreen reporter (p65^NGreen^) and stained with anti-p65 fluorescent labeled antibody (p65^Ab^) with no treatment (“NT”, top) or 40 min after TNF*α* treatment (bottom). Scale bar, 100µm. **c.** InCITE antibody detects p65 nuclear translocation. Flow cytometry measured distribution of HeLa nuclei stained with p65^inCITE Ab^ (followed by secondary antibody stain conjugated with Alexa Fluor 647) with no treatment (“NT”, blue) or 40 min after TNF*α* treatment (red). **d.** InCITE-seq detects p65 nuclear translocation. Distribution of p65 levels (nCLRs) in NT (blue) and TNF*α* treated (red) nuclei profiled by inCITE-Seq (*P*<0.0001, t-test). **e,f.** InCITE-seq reveals p65 expression program. **e.** Expression (Z score, color bar) of top genes (rows) positively associated with p65 levels by a linear model (top, **Methods**) across the cells (columns), visualized for the top decile (p65^high^) and bottom decile (p65^low^) of p65 nuclear protein levels by inCITE-seq (bar plot, top, CLR). **f.** Enrichment (-log_10_(P-value), x axis, hypergeometric test) of Gene Ontology terms (y axis) enriched in 142 genes significantly positively associated with p65 levels.

## Results

### InCITE-seq detects nuclear translocation of a TF induced by an extracellular signal

We first developed inCITE-seq in a cell line to detect cell state changes, defined as changing nuclear levels of a transcription factor that translocates into the nucleus in response to an external stimulus. We used a HeLa cell line expressing a p65-mNeonGreen reporter construct^47^, where p65 is localized to the cytoplasm in untreated cells and translocates to the nucleus upon TNF*α* stimulation (**Fig. 1b**; **Methods**). At peak nuclear translocation (∼40 min post-stimulation^48^), total p65 levels in the whole cell are constant, with no discernible difference between untreated and TNF*α* stimulated cells in mNeonGreen signal measured by flow cytometry, but nuclear p65 levels are highly elevated (**Supplementary Fig. 1a,b**). Using a DNA-conjugated anti-p65 antibody (p65^inCITE-Ab^), we optimized intranuclear staining conditions, such that flow cytometry successfully resolved p65 levels between nuclei from untreated *vs*. TNF*α* stimulated cells (**Fig. 1c**; **Supplementary Fig. 1c**; **Methods**). We then profiled 10,014 single nuclei from untreated and TNFα treated unsorted populations stained with p65^inCITE-Ab^ and barcoded by nucleus hashing^46^ using inCITE-seq (**Methods**). We estimated antibody levels as counts of antibody-derived tags (ADTs)^8^ normalized by the counts of the nucleus hashtag (HTO) to yield nuclear ADT (“nADT”) units to account for differences in poly-dT capture on beads (**Supplementary Fig. 1d**), and log-scaled to nuclear centered log ratios (nCLRs)^8, 49^ (**Methods**). The distribution of p65 levels differed significantly between untreated and TNF *α* stimulated nuclei (**Fig. 1d;** *P<*10^-4^, t-test). The quality of the associated snRNA-seq profiles was comparable to other snRNA-seq data from human-derived cell lines, as reflected by the number of unique transcripts (UMIs) and genes, with a slight reduction in library complexity, which we ascribe to formaldehyde fixation^50^ (**Supplementary Fig. 1e**).

Genes whose RNA levels correlated with nuclear p65 protein levels were known targets of NF-*κ*B, confirming that inCITE-seq captured the expected protein level changes in activated cell states. Specifically, to identify genes whose expression was associated with p65 levels we used a linear model, fitting each gene’s expression as a function of p65 levels, controlling for cell cycle (**Methods**). The 142 genes positively associated with p65 levels (at FDR 1%; **Methods**) were enriched for pathways such as response to cytokine (**Fig. 1f**, P<3*10^-9^, hypergeometric test, **Methods**), and included well-known NF-*κ*B targets, such as *CXCL8*, *NFKBIA*, and *TNFAIP3* (**Fig. 1e**). Notably, and as expected^15^, RNA and protein levels of p65/*RELA* were not well-correlated at this time scale of stimulation (Pearson *r*^2^=0.0008, *P*=0.004) (**Supplementary Fig. 1f**). Taken together, these results show that inCITE-seq accurately quantifies nuclear protein and RNA levels, which can be integrated to identify putative targets of a TF.

### Profiling the mouse hippocampus with inCITE-seq after pharmacological treatment

Next, we turned to an *in vivo* setting and applied inCITE-seq to profile the mouse hippocampus after treatment with kainic acid (KA), a potent agonist of glutamate receptors that induces seizure, involving multiple cell types and pathways resulting in neuronal death, neuroinflammation, and oxidative stress^51–53^. As the complexes NF-kB and AP-1 are involved in neuroinflammation, synaptic remodeling, and signal transduction of glutamate receptors^51, 54–56^, we asked how their components p65 and c-Fos are linked to gene expression changes during early response to seizure. We profiled nuclei from the hippocampus two hours after KA treatment with multiplexed protein measurements of p65, c-Fos, the neuronal marker and regulator of alternative splicing NeuN^57^, and the TF and microglial lineage marker PU.1^58^ (**Fig. 2a**; **Methods**). Flow cytometry verified that intranuclear staining with NeuN and PU.1 captured cell subsets at the expected proportions (58.3% and 2.9% of all nuclei, respectively; **Supplementary Fig. 2a,b**); NeuN fluorescence levels and side scatter were directly correlated, consistent with reports that NeuN levels are elevated in enlarged nuclei with decondensed chromatin^59, 60^. The proportion of c-Fos^high^ increased as expected in response to KA, from 0.21% to 48.7% (**Supplementary Fig. 2d,e**).

**Figure 2.**
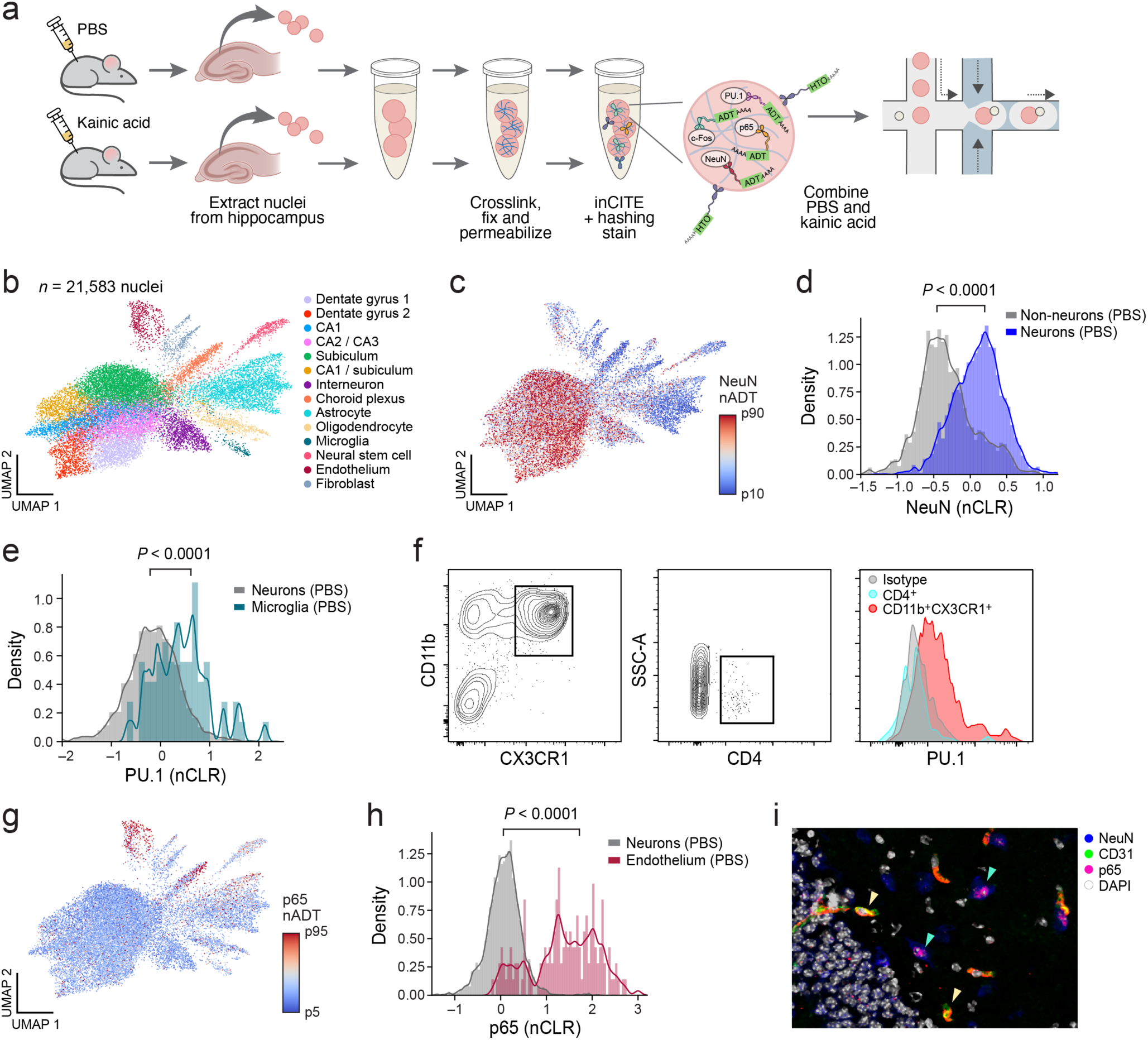
*In vivo* application of inCITE-seq shows cell type-specific protein expression in the mouse hippocampus. **a.** InCITE-seq of the mouse hippocampus after kainic acid or PBS (control) treatment, with nucleus hashing. **b.** Major cell types from the adult mouse hippocampus identified from inCITE-seq. UMAP embedding of 21,583 snRNA-seq profiles (dots) from inCITE-seq colored by cluster and annotated *post hoc* (color legend). DG, CA, and subiculum clusters refer to excitatory neurons from these regions. **c,d.** Higher NeuN expression in neurons by inCITE-seq. **c.** UMAP embedding (as in (**b**)) of inCITE-seq nuclei RNA profiles colored by NeuN protein levels (nADT, Color scale from 10th to 90th percentile). **d.** Distribution of NeuN protein levels (nCLR, x axis) in neurons (blue) and non-neural (gray) nuclei from one batch of PBS treated mice. Curve: kernel density estimate. *P*<0.0001, t-test. **e,f.** Increased PU.1 levels in microglia. **e.** Distribution of PU.1 protein levels (nCLR, x axis) in microglia (turquise) and neurons (gray) from one batch of PBS treated mice. Curve: kernel density estimate. *P*<0.0001, t-test. **f.** Right: Distribution of PU.1 in microglia (CD11b^+^ CX3CR1^+^, red), CD4^+^ cells (blue) and isotype (gray) cells measured by flow cytometry (left and middle panels). **g-i.** p65 is enriched in the endothelium and subpopulations of the choroid plexus. **g.** UMAP embedding (as in (**b**)) of inCITE-seq nuclei RNA profiles colored by p65 protein levels (nADT, color scale from 5th to 95th percentile). **h.** Distribution of p65 levels (nCLR, x axis) in endothelial (red) and neurons (gray) nuclei (*P*<0.0001, t-test). **i.** Immunofluorescence stain of the hippocampus with endothelial marker CD31 (green), NeuN (blue), and p65 (pink). Yellow arrowheads: co-localization of CD31 and p65. Green arrowheads: lowly expressed p65 in neurons.

We profiled 21,583 nuclei with inCITE-seq across control (PBS) and KA treated mice, capturing all major cell types of the hippocampus. Using only the snRNA-seq profiles, we identified cell types by unsupervised clustering after addressing ambient RNA^61^, batch-correction, and regressing out treatment (**Supplementary Fig. 3a,b**; **Methods**), followed by *post hoc* annotation with hippocampal cell type specific markers^62^. All major cell types of the hippocampus formed discernible clusters, including excitatory neurons from distinct anatomical regions of the hippocampus (dentate gyrus, CA1, CA2/3, and subiculum), astrocytes, and neural stem cells (**Fig. 2b**; **Supplementary Fig. 3c**). Notably, the separation between clusters was not as strong as in snRNA-seq of the hippocampus^2^; if future studies employ inCITE-seq in tissues without a reference atlas, jointly embedding the RNA profiles from regular and fixed snRNA-seq or inCITE-seq should help provide discrete clusters for annotation^11^.

Nuclear protein levels measured by inCITE-seq reflected expected cell type-specific differences based on RNA-defined clusters at baseline (PBS control). The neuronal marker NeuN was elevated in neuronal clusters (dentate gyrus clusters, CA clusters, subiculum clusters, interneuron) as expected (**Fig. 2c,d**; *P*<10^-4^, t-test two-sided; **Supplementary Fig. 4a, Methods**); low NeuN in subpopulations of astrocytes may reflect signal leakage, as NeuN is abundant in this tissue. PU.1, the microglial marker and lineage-specifying TF, was significantly higher in microglia nuclei compared to neurons (**Fig. 2e**; *P*<10^-4^, t-test, two-sided); the lower signal-to-noise ratio of PU.1 could reflect excess antibodies targeting a small population, which could be improved with antibody titrations^63^. Using flow cytometry, we confirmed that PU.1 was microglia-specific by co-staining with microglia cell surface markers CD11b and CX3CR1, showing high levels of PU.1 in CD11b^high^CX3CR1^high^ populations but not in CD4^high^ cells (**Fig. 2f**). Nuclear expression of p65 was highly enriched in endothelial nuclei, consistent with expression of its encoding gene *Rela* in this population (**Fig. 2g,h**; **Supplementary Fig. 3d**), and in subsets of nuclei from the choroid plexus (**Supplementary Fig. 5a**). Immunohistochemistry of p65 with the endothelial marker CD31 and neuronal marker NeuN verified high levels of p65 expression in the endothelium (**Fig. 2i**, yellow arrowheads), and lower levels in neurons^64^ (**Fig. 2i**, green arrowheads).

### InCITE-seq captures upregulation of activity-induced c-Fos expression in neurons

KA treatment induced c-Fos expression in neuronal nuclei, with variations across neuronal subtypes. InCITE-seq measures showed widespread expression of nuclear c-Fos across multiple cell types (**Fig. 3a**), with significant upregulation in neuronal nuclei of KA treated mice compared to PBS (**Fig. 3b**; *P*<10^-4^, t-test, two-sided). Interestingly, subsets of neuronal nuclei differed in c-Fos levels, such that neuronal nuclei from CA had lower levels than those from either DG or interneuron, which were comparably high (**Fig. 3c**, **Supplementary Fig. 5b,c**; *P<*0.0001 and *P=*0.0009, t-test, two-sided). In contrast, p65 nuclear levels did not change from KA treatment at this time point (**Fig. 3d**), consistent with previous reports^56, 65^. These patterns were confirmed by immunofluorescence, where c-Fos is expressed in the soma of multiple neuronal types, including the dentate gyrus, CA neurons, and interneurons, with higher c-Fos intensity in DG and somatostatin (SST+) interneurons (IN) compared to CA neurons (**Fig. 3e,f**), and no change in p65 due to treatment (**Fig. 3g**). Overall, inCITE-seq quantitatively measured nuclear protein levels that reflected diverse levels of activity-regulated TFs across cell types.

**Figure 3.**
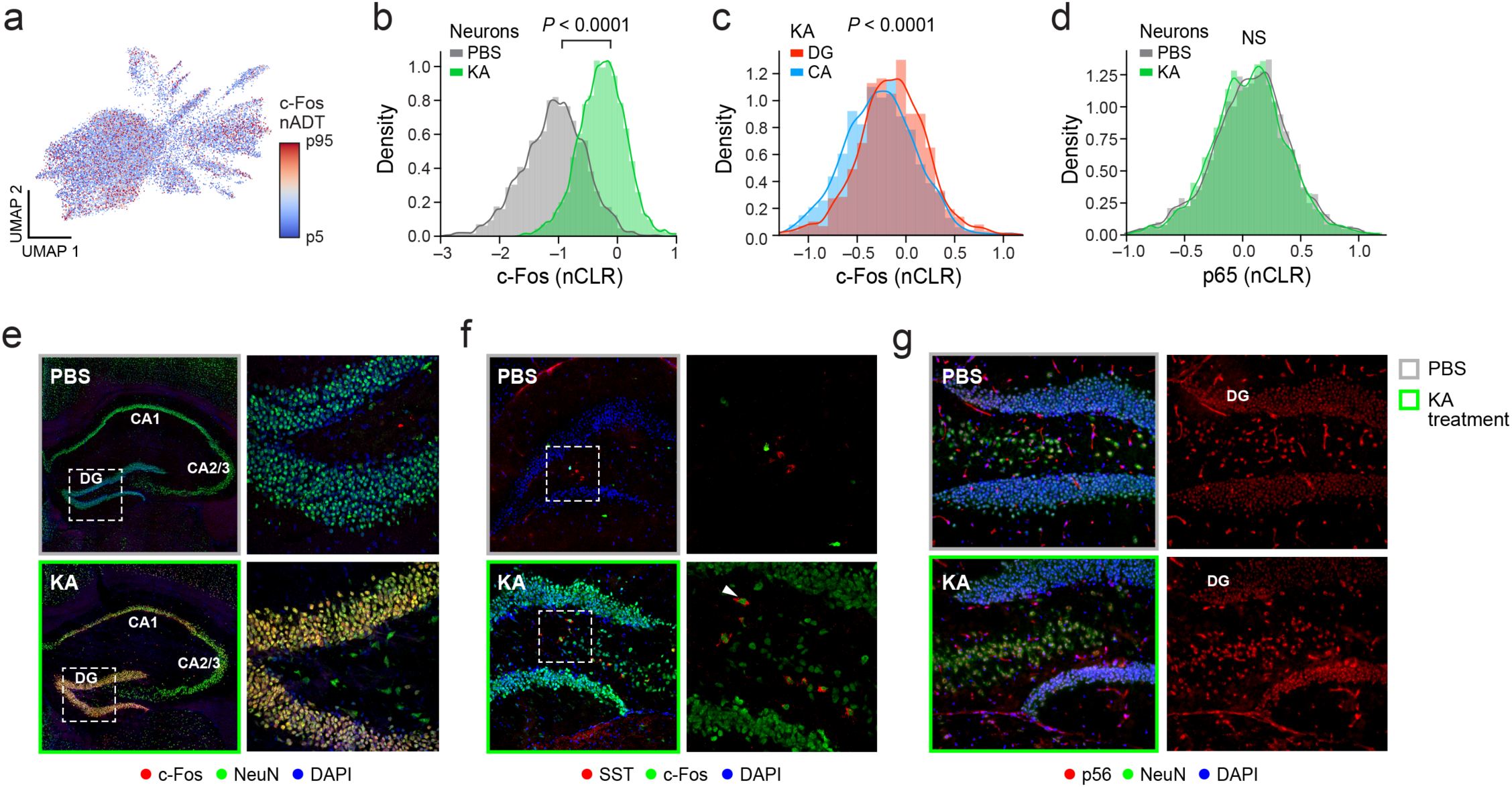
InCITE-seq measures changes in nuclear TF levels after stimulation of the mouse hippocampus. **a.** Widespread expression of c-Fos across nuclei subsets. UMAP embedding (as in Fig. 2b) of inCITE-seq nuclei RNA profiles colored by c-Fos protein levels (nADT, color scale from 5th to 95th percentile). **b-e.** Increased c-Fos levels, especially in dentate gyrus neuronal nuclei, following KA treatment. Distribution of c-Fos (**b,c**) or p65 (**d**) protein levels (nCLR, x axis) and a kernel density estimate (curve) from single batches in neurons of KA vs. PBS treated mice (**b**, *P*<0.0001, t-test), dentate gyrus (DG) vs. cornu ammonis (CA) neurons in KA treated mice (**c,** *P*<0.0001, t-test), or neurons of KA vs. PBS treated mice (non-significant; *P*=0.063, t-test). **e.** Immunofluorescence stain of c-Fos (red), NeuN (green), DAPI (blue) in the hippocampus after PBS (gray border) or KA (green border) treatment. Major hippocampal features denoted: dentate gyrus (DG) and cornu ammonis (CA). Left: full image. Right: zoom image of the DG (dashed area on left), showing heterogeneity in c-Fos intensity. **f.** Increased c-Fos expression in SST^+^ hippocampal interneurons after KA treatment. Immunofluorescence stains of somatostatin (SST, red), c-Fos (green), and DAPI (blue) in the hippocampus after PBS (gray border) or KA (green border) treatment. Right: zoom image of the DG (dashed area on left). **g.** p65 is widely and lowly expressed in neurons under both PBS and KA treatment. Immunofluorescence stains of p65 (red), NeuN (green), and DAPI (blue) in in the hippocampus after PBS (gray border) or KA (green border) treatment.

### Modeling genes associated with each protein recovers known TF targets

Next, we devised an approach to infer the impact of each nuclear TF on gene expression. To elucidate the global impacts of TFs, we first regressed out cell type (cluster), treatment, and their interaction to account for collinearity, especially between treatment and c-Fos levels; we then modeled the residuals of each gene as a linear combination of c-Fos, p65, PU.1 and NeuN^66^ (**Methods**). Genes significantly associated with the three TFs were interpreted as putative TF-regulated genes, with the effect size of each TF estimated by their coefficient. TF genes comprised known targets and pathways (**Fig. 4a-c**; **Supplementary Fig. 6b**), with p65 predicted to impact for example autotaxin (*Enpp2*), which is induced by overexpression of p65 *in vitro*^67^, and PU.1 predicted to regulate known microglia markers and targets, such as *Trem2*, *Tyrobp* and *C1qa*^68, 69^. c-Fos associated genes *Npas4, Nr4a1*, and *Junb* reflected broad activity-induced upregulation^37, 42^, as well as upregulation of its own transcript *Fos*. Although our model also identified NeuN-associated genes (**Supplementary Fig. 6a**), we reasoned that these genes could reflect either direct changes in transcript levels via splicing or indirect effects of a generally active transcription state, as NeuN is associated with decondensed chromatin and enlarged nuclei^59, 60^.

**Figure 4.**
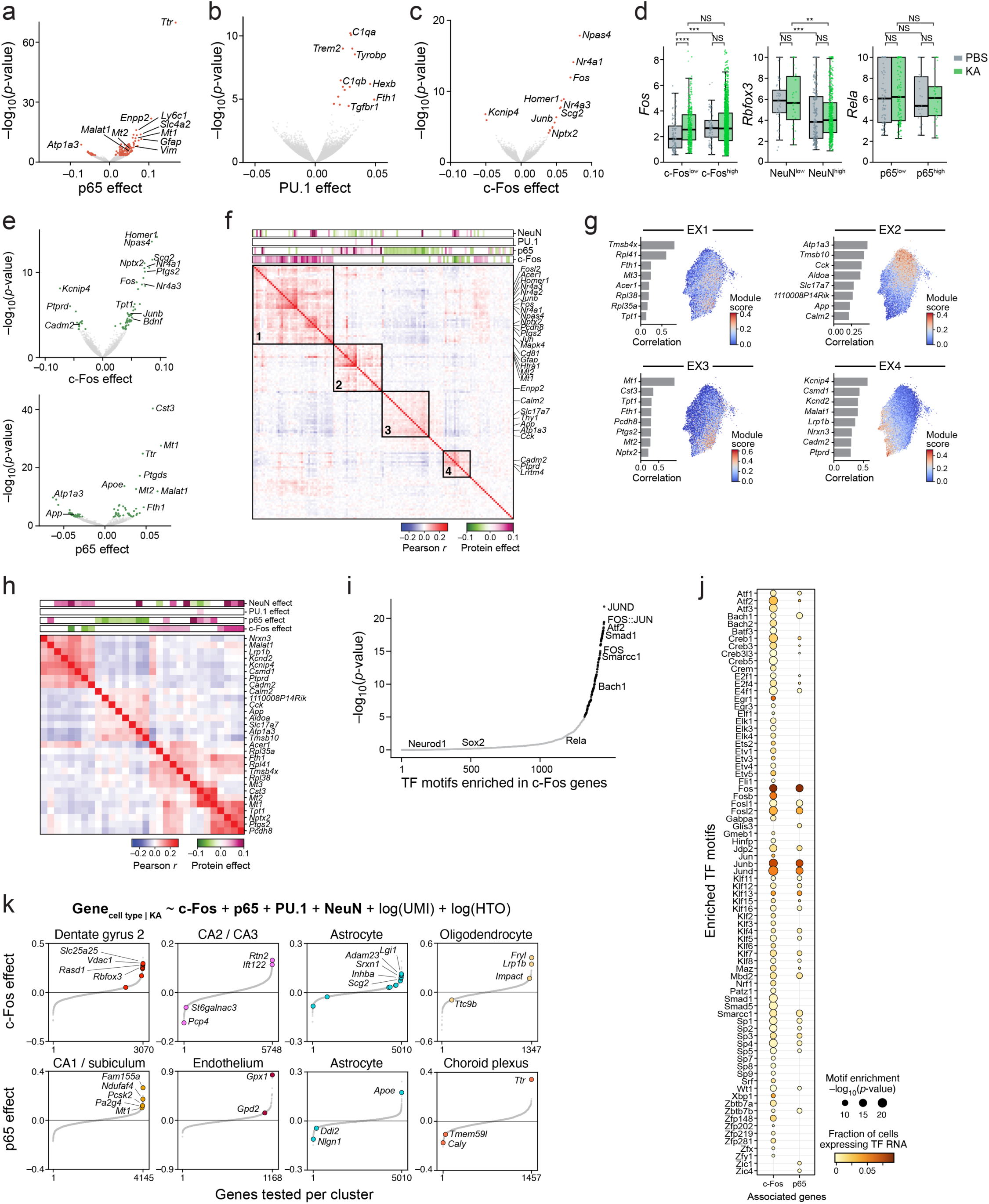
Inferring TF effects on gene and module expression from inCITE-seq data. **a-c.** Global association of TFs to genes. Significance (y axis, -log_10_(P-value)) and effect size (x axis) for genes (dots) globally associated with p65 (**a**), PU.1 (**b**), and c-Fos (**c**) by a model of gene expression as a linear combination after regressing out treatment and cell type (**Methods**). Select genes are labeled. Colored dots: Benjamini-Hochberg FDR<5%. **d.** Relation between RNA and nuclear protein levels of each TF. Distribution of RNA levels (Z score of log-normalized counts, y axis) in nuclei with high or low levels (**Supplementary Fig. 4**) of the encoded protein (x axis) in PBS (gray) or KA (green) treatment. Boxplot shows the median, with ends representing 25% and 75% quartiles and whiskers at 1.5 interquartile range below 25% or above 75% quartiles. Dots: individual nuclei with non-zero mRNA levels. **P<10^-2^, ***P<10^-3^, **** *P*<10^-4^, t-test. NS – not significant. **e.** Genes associated with each TF in excitatory neurons. Significance (y axis, -log_10_(P-value)) and effect size (x axis) for genes (dots) globally associated with c-Fos (left) or p65 (right) by a model of gene expression as a linear combination after regressing out treatment (**Methods**). Select genes are labeled. Colored dots: Benjamini-Hochberg FDR<5%. **f.** Pearson correlation coefficient (red/blue colorbar) of gene expression across EX neurons (rows and columns) significantly (FDR<5%), for genes associated positively (purple) or negatively (green) with either c-Fos or p65 within EX neurons, ordered by hierarchical clustering. Top bars: Effect size of each protein from the linear model (**Methods**). Black boxes 1-4: modules. **g,h.** Association of NMF- derived gene programs in EX neurons with TFs. **g.** Right: UMAP embedding of EX neuron nuclei RNA profiles colored by the NMF module score. Left: Top 8 module genes (y axis) and their Pearson correlation with module scores (x axis). **h.** Pearson correlation coefficient of gene expression between the top 8 genes of the EX neuron NMF programs (rows), ordered by hierarchical clustering. Top bars (purple/green): significant effect size of each protein from the linear model. **i,j.** Prediction of co-regulatory patterns by TF motif enrichment. **i.** Significance (- log_10_(P value), y axis, and x axis rank order) of TF motif (dots) enrichment in enhancers of genes associated with c-Fos in EX neurons. Black: significantly enriched motifs (*P<*10^-5^, hypergeometric test); Gray: not significant motifs (two lineage-specifying TFs labeled). **j.** TF motif enrichment (dot size, -log_10_(P-value)) and proportion of EX neuron nuclei expressing the TF’s RNA (color) of TFs (columns) whose motifs are significantly enriched in enhancers of c-Fos or p65 (rows) associated genes. **k.** Effect size (y axis, with rank order on x axis) of association with c-Fos (top) or p65 (bottom) in specific cell types (labeled on top) after KA treatment (**Methods**). Color: significantly associated genes in color (Benjamini-Hochberg FDR<5%).

### Relating protein and mRNA levels of inCITE target genes

Comparing nuclear protein levels with levels of their own encoding mRNA revealed a wide range of regulatory dynamics. We partitioned nuclei by protein levels into “high” and “low” bins (with batch-specific thresholds, **Supplementary Fig. 4**) and compared the cognate mRNA level between bins (**Fig. 4d**). *Fos* mRNA was higher in c-Fos^high^ compared to c-Fos^low^ (*P=*0.0015, t-test), but only under PBS treatment, whereas in KA treated populations, *Fos* mRNA was highly expressed regardless of c-Fos protein levels. This is consistent with a model where c-Fos regulates *Fos* expression, but treatment had saturated *Fos* induction^31^. In contrast, NeuN and the mRNA of its encoding transcript *Rbfox3* were inversely correlated (*P<*0.01, t-test), although unspliced, intron-retaining pre-mRNA levels of *Rbfox3* (estimated by Velocyto^17^, **Methods**) were upregulated by KA (**Supplementary Fig. 6c**). This is consistent with observations for other RNA-binding proteins that negatively regulate their own transcript expression^70, 71^. *Rela* mRNA levels did not differ significantly between p65^high^ and p65^low^ nuclei (*P=*0.78, t-test), and *Spi1* mRNA transcripts encoding PU.1 were not detected, underscoring the importance of protein measurements as a complement to RNA, especially for lowly expressed TFs^72^.

### Genes associated with each TF are co-expressed as distinct modules within cell types

Next, we probed how the expression of TF associated genes relate to each other by examining their co-expression patterns within cell types. We again modeled each gene within excitatory neurons (EX; dentate gyrus, CA, and subiculum clusters) and astroglia (astrocytes and microglia) as a linear combination of the four proteins after regressing out treatment (**Methods**; **Fig. 4e; Supplementary Fig. 7a,b**). We then clustered genes associated with c-Fos and p65 based on their co-expression patterns, revealing distinct gene modules (**Fig. 4f; Supplementary Fig. 7c**). Interestingly, overlaying the effect of protein levels revealed that each module reflected a unique set of protein effects (**Fig. 4f**). Specifically, within excitatory neurons, genes in module 1 are positively associated with c-Fos protein levels in the nucleus, while modules 2 and 3 are respectively positively and negatively regulated associated with p65 levels, and module 4 reflected a mixture of both c-Fos and p65 levels. Our results show that we can attribute TF levels to distinct sets of genes, and disentangle gene modules that are upregulated or downregulated by the same TF.

The coupling between co-expressed genes and TF effect types led us to ask whether gene programs that are normally identified from expression alone^73, 74^ also reflected protein effects. Using non-negative matrix factorization on gene expression, we independently identified gene programs within each cell type (4 programs in EX neurons, 3 programs in astroglia; **Fig. 4g; Supplementary Fig. 7d; Methods**). Some of these programs were upregulated in response to treatment (**Supplementary Fig. 7e,f**). We examined the co-expression of the top 10 genes of each program by clustering, then overlaid the effect of protein levels, which showed that each program was also specific TF regulatory effects (**Fig. 4h, Supplementary Fig. 7g**). Altogether, our approach expands beyond the identification of gene expression modules, allowing the identification of specific TF contributions to the regulation of gene expression, and even disentangling contributions from multiple TFs, which can increase the interpretability of gene expression programs.

We next predicted other TFs that could co-regulate genes associated with c-Fos and p65 within excitatory neurons, by testing for *cis* regulatory motifs enriched in the enhancer regions of the TF-associated genes using differentially accessible regions (DARs) of the mouse hippocampus previously profiled in saline- and KA- treated mice (1 hr post treatment)^75^ (**Methods**). TF motifs in enhancers of c-Fos and p65- associated genes belonged to components of the AP-1 complex and related TFs, including Jun, JunD, and Atf2 (**Fig. 4i**), which is a mediator of excitotoxicity in KA-induced seizure models, and is phosphorylated by c-Jun^76^. Other TFs included *Egr1* and *Egr3*, consistent with direct effects of c-Fos on chromatin following KA treatment^77, 78^, and members of the Elk, Etv, and Klf/Sp families, all lowly expressed at the RNA level in our data (**Fig. 4j**). TF motifs enriched in KA-treatment DARs of c-Fos and p65 associated genes *vs*. non-associated genes further underscored the potential role of Klf/Sp families (**Supplementary Fig. 8a**). Notably, motifs of NF-κB components (Rel, Rela, Relb, Nfkb1, Nfkb2) were not significantly enriched. This suggests that p65-associated genes may reflect indirect effects of p65, and that direct effects may occur at a different time scale, or depend on the direction of the TF effect (*i.e.* downregulated vs. upregulated).

### c-Fos and p65 may regulate genes involved in oxidative stress after seizure

Finally, we analyzed how treatment impacted TF effects on individual genes at a global scale and within individual cell types. To assess the global impact, we modeled RNA levels as a linear combination of the four proteins separately within each treatment (PBS or KA) and compared each TF effect on genes across the treatments (**Methods**; **Supplementary Fig. 8b**). The impact of p65 on gene expression were largely consistent across treatment, as reflected by correlated effect sizes between PBS and KA (R^2^=0.11, *P*=6*10^-263^). In contrast, c-Fos effects varied by treatment as reflected by no correlation of the effect sizes between PBS and KA (R^2^=8*10^-6^, *P*=0.77). Genes that were specifically associated with c-Fos after KA treatment included *Ptgs2* (encoding cyclooxygenase-2, COX-2), a responder to oxidative stress after traumatic brain injury^79–81^. We further examined TF effects after KA treatment at a cell type-specific level (**Methods**; **Fig. 4k**). Genes associated with c-Fos included mitochondria-associated genes *Slc25a25* and *Vdac1* in dentate gyrus, *Lgi1* and *Adam23* in astrocytes, and *Lrp1b* in oligodendrocytes. *Lgi1* causes an inherited form of epilepsy^82, 83^, and encodes a secreted protein that modulates synaptic strength by forming a ligand-receptor pair with *Adam22* and *Adam23*^84, 85^. While their interaction in epilepsy is well-appreciated for neurons, how activity-dependent astrocytic expression of these two genes modulates circuits remains to be seen. Interestingly, the antioxidant enzyme glutathione peroxidase 1 (*Gpx1*) increased with p65 in endothelial cells. Mice overexpressing *Gpx1* were protected against oxidative stress and ischemic cell death^86^, and overexpression of p65 in a kidney-derived cell line led to *Gpx1* expression^87^. As seizures lead to reactive oxygen species (ROS) that activate the NF-κB pathway^88–90^, our results suggest how p65 may regulate genes that protect against oxidative stress in a cell type-specific manner. Overall, these results show that the impact of TFs on individual genes could change depending on context.

## Discussion

In conclusion, inCITE-seq reliably profiles protein and RNA levels in nuclei, opening the way to better understand the relationship between TFs and their targets, including during dynamic responses, in the native context of tissues. Nucleus-based multimodal profiling surmounts key technical challenges, and enables characterization of cells from tissues that either are hard to dissociate or are archived in frozen form, especially clinical specimens from human disease studies, including cancer and neurodegeneration. Furthermore, our approach could be easily adapted to enrich for snRNA-seq of subpopulations^91^. We note that one drawback of inCITE-seq and related approaches is the need for high-quality antibodies. Future studies can combine inCITE-seq with metabolic labeling^50, 92–94^, antibodies targeting phosphorylated forms of TFs, joint RNA and chromatin accessibility profiles^95^ combined with spatial inference^96^, and samples collected across time, for refined understanding of gene regulation in tissues. By measuring multiple TFs simultaneously, inCITE-seq opens the way to decipher complex phenotypes and regulatory mechanisms in development, when TF combinations are key to define cell type diversity^97^, or in behavior and learning, and help recover the impact of ligands acting on different and multiple targets across cell types, and in disease, where GWAS highlights the role of variants in regulatory regions.

## METHODS

### Mice

C57BL/6J (Jax 000664) mice were purchased from The Jackson Laboratory and bred in-house. Male mice were used at ∼8 weeks of age. All mice were maintained under SPF conditions on a 12-h light–dark cycle, and provided food and water ad libitum. All mouse experiments were approved by, and performed in accordance with, the Institutional Animal Care and Use Committee guidelines at Weill Cornell Medicine.

### Cell culture

HeLa cells expressing a p65-mNeonGreen reporter construct (gift from Jonathan Schmid-Burgk)^47^ were cultured at 37°C and 5% CO_2_ in DMEM with high glucose, pyruvate, GlutaMax^TM^ (ThermoFisher Scientific 10569010), heat-inactivated fetal bovine serum (ThermoFisher Scientific 16000044) and 100 U/mL penicillin-streptomycin (ThermoFisher Scientific 15140163). For immunohistochemistry to visualize nuclear translocation in whole cells, cells were seeded in 6 well plates containing poly-L-lysine treated #1.5 glass coverslips (Thomas Scientific 1217N81) at a density of ∼5×10^4^ cells/mL 24 hours prior to TNF*α* stimulation. For inCITE-seq, HeLa cells were seeded in 10cm Petri dishes at least 24 hours prior to TNF*α* stimulation. Cells reached 70-80% confluence prior to stimulation and extraction.

### Stimulation of HeLa cells with TNF*α*

HeLa cells were stimulated with 30 ng/mL of TNF*α* at 37°C for 40 min to induce translocation of p65 into the nucleus. Media was removed and HeLa cells were washed with 1X PBS. HeLa cells for nucleus isolation were scraped in 2mL of EZ lysis buffer (Sigma Aldrich N3408) and moved to a 15mL falcon tube for nucleus extraction.

### Kainic acid injection of mice

8-week-old male mice were first acclimated into the procedure room for 1h prior to seizure induction. Mice were injected *i.p.* with either PBS or 20mg/kg kainic acid (KA; Sigma K0250) dissolved in PBS. All animals were observed continuously for two hours and scored using a modified Racine scale^98^: stage 0, normal behavior; stage 1, immobility and rigidity; stage 2, head bobbing; stage 3, forelimb clonus and rearing; stage 4, continuous rearing and falling; stage 5, clonic-tonic seizure; stage 6, death. Any mice not reaching at least stage 1 by 30 minutes post-KA were given an additional injection of 10mg/kg KA to facilitate seizure activity. After 2 hours, mice were euthanized by CO_2_, perfused through the left ventricle with 20mL of PBS, and the entire brain was removed. The hippocampus was then dissected on ice before freezing on dry ice.

### Immunohistochemistry of HeLa cells

Coverslips (#1.5, 18mm, Thomas Scientific 1217N81) treated with poly-L-lysine solution seeded with HeLa cells and grown in 12-well tissue culture plates were stimulated with TNF*α* as described. Wells were washed with PBS, fixed with 4% PFA at RT for 15 min, then washed 3x with PBS. All staining was on Parafilm, with cells facing down, such that solution volumes (100µL) were sandwiched between the coverslip and Parafilm. Cells were blocked at RT for 30 min (1X PBS, 5% normal goat serum, 0.3% Triton X-100), incubated with 1:200 p65^Ab^ (BioLegend cat #622601) in 5% BSA and 0.02% Tween 20 for 1 hour at RT, and washed 3x with PBST (1X PBS, 0.02% Tween 20). Cells were then treated with anti-rabbit Alexa Fluor 647 secondary (Invitrogen A27040) at 1:1,000 in PBST for 1hr in dark, RT, then washed 4x with PBST, with the final wash containing 1:1,000 DAPI. Coverslips were mounted onto SuperFrost slides (Fisher Scientific 22-037-246) with antifade (ThermoFisher Scientific S36937) and sealed with nail polish. Slides were stored at 4°C until ready for imaging.

### Immunohistochemistry of mouse hippocampus

Mice were euthanized 2 hours after PBS or KA administration and perfused through the left ventricle with 20mL of PBS before gently removing the entire CNS. Brain tissue was then immediately immersed in OCT medium and quickly frozen in dry ice, followed by sectioning at 10 µm using a cryotome and collection on charged slides which were then frozen on dry ice and stored at -30°C until further use. Slides were removed from storage and fixed with 4% PFA for 10 minutes at room temperature, washed once in PBS, then blocked with PBS containing 0.1% Triton-X 100, 5% normal donkey serum and 5% normal goat serum (Jackson Immunoresearch) for 30 min at room temperature. Sections were incubated with the following primary antibodies at the indicated dilution in blocking buffer overnight at 4°C: NeuN (BioLegend 1B7, cat #834502), p65 (BioLegend Poly6226, cat #622601), c-Fos (BioLegend Poly6414, cat #641401), CD31 (eBioscience 390, cat #14-0311-82). Sections were then washed in PBS 3 times before incubating in species specific anti-IgG secondary antibodies conjugated to AF488, AF594, or AF647 (Jackson Immunoresearch) diluted 1:500 in blocking buffer for 1 hr at room temperature. Sections were then washed once in PBS, once in PBS containing DAPI, and a final time in PBS before mounting (Prolong Diamond Antifade, ThermoFisher). Slides covered with coverslip were dried overnight, sealed with clear nail polish, and imaged.

### Microscopy

HeLa cells and mouse hippocampal sections were imaged on an Olympus Fluoview FV1200 biological confocal scanning microscope at 20X or 40X (Olympus, LUCPLFLN) with sequential laser emission and Kalman filtering. Images were processed with ImageJ.

### FACS analysis of PU.1 stain in microglia suspensions

Mice were euthanized with CO_2_ and perfused through the left ventricle with 20mL of PBS. Whole brain was removed and placed in 2.5 mL of digestion buffer (PBS, 5% FCS, 1mM HEPES) before finely chopping. 400U of Collagenase D (Roche) were then added to the mixture, which was incubated at 37°C for 30 min before the addition of 50 μL of 0.5M EDTA and an additional 5 min incubation. Digested tissue was then mashed through a 40µm cell strainer, pelleted at 700g in a swinging bucket centrifuge, then resuspended in 10mL of 38% isotonic Percoll, and centrifuged at 2000 RPM for 30 min with no brake. The myelin debris layer was removed by aspiration and the cell pellet washed once in PBS. Cells were then blocked with 1:100 FcX (BioLegend 156604) before a 15 min incubation with the following antibodies at a 1:200 dilution in PBS: CD45.2-FITC (eBioscience 104 cat #11-0454-82), CD4-BUV395 (BD GK1.5 cat #563790), CD11b-BV421 (BioLegend M1/70 cat# 101235, CX3CR1-APC (BioLegend SA011F11 cat #149008). Cells were washed once in PBS and then fixed and permeabilized using the Foxp3/Transcription Factor Staining Buffer Set (eBioscience) before staining with PU.1-PE (BioLegend 7C2C34 cat #681307) or rat IgG2a-PE isotype (BioLegend RTK2758, cat #400507) for 30 minutes at room temperature. Cells were washed once, resuspended, run on a LSRFortessa cytometer (BD Biosciences) and analyzed with FlowJo software (Tree Star).

### Nucleus extraction

Nuclei from tissue or cell lines were extracted using EZ Prep (Sigma-Aldrich N3408) and Glass Dounce Kit (Sigma Aldrich D8938) as previously described^2^. Briefly, tissue or cells were placed in 2mL of EZ lysis buffer containing Recombinant RNase Inhibitor (Takara Bio 2313A), dounced 24 times with pestle A, and dounced 24 times with pestle B. Nucleus solutions were transferred to a 15mL Falcon tube. An additional 3mL of EZ lysis buffer was used to wash out the glass mortar and added to the 15mL Falcon tube. Solutions were incubated on ice for 5 min, pelleted with a swinging bucket centrifuge at 500g for 5 min at 4°C, resuspended in 5mL EZ lysis buffer with a P1000 pipette, incubated on ice for 5 min, and pelleted as in previous step. Nuclei were resuspended in 1mL pre-chilled buffer (1X PBS, 3mM MgCl2, Recombinant RNase Inhibitor at 40 U/mL (Takara Bio 2313A)) and filtered through a 35µm FACS tube (Falcon 352235). Tris-based buffers were avoided to maintain the efficacy of formaldehyde fixation by preventing the excess free amines in Tris from quenching fixation.

### Intranuclear antibody stain of nucleus suspensions

Nuclei were simultaneously fixed and permeabilized by adding 3mL of 1.33% FA-NT (1.33% formaldehyde, 0.2% NP-40, 0.1% Tween 20, 3µL glacial acetic acid) to 1mL of nuclei suspended in PBS with 3mM MgCl_2_. Samples were incubated for 10 min at 4°C with rocking. The fixation reaction was quenched by spiking in 3µL of 1M glycine. Nuclei were filtered through a 20µm strainer (pluriSelect 431002040) and pelleted in a swinging bucket centrifuge at 850g, 5 min, 4°C, then resuspended in 500µL of blocking buffer (1:100 FcX (BioLegend 156604) + 1% UltraPure BSA (ThermoFisher Scientific AM2618) + 0.05% Dextran Sulfate), incubated for 15 min at 4°C with rocking, and pelleted as previously described. Pellets were resuspended in 200µL of primary antibodies diluted in blocking buffer solution and incubated at 4°C for 1 hour with rocking. Primary antibody concentrations were as follows: p65^Ab^ (raised in rabbit) at 1:400, p65^inCITE Ab^ (raised in rabbit) at 1:400, c-Fos^inCITE Ab^ (raised in rabbit) at 1:400, NeuN^inCITE-Ab^ (raised in mouse) at 1:500, PU.1^inCITE Ab^ (raised in rat) at 1:200. Nucleus hashing antibodies (BioLegend cat #682213, 682215) were simultaneously added to each sample at 1:200. After incubation, nuclei were pelleted using a swinging bucket centrifuge at 850g for 5 min at 4°C (centrifuge conditions for all subsequent spins), washed 2x with 500µL of 0.2% PBST, incubated for 5 min, and re-pelleted. Nuclei were then either resuspended in 300µL of 1X PBS in preparation of loading on the 10x Genomics Chromium instrument (below), or resuspended in 200µL of secondary antibodies at 1:1000 dilution in blocking buffer and 10x DAPI, incubated in the dark at 4°C for 30 min, washed twice as previously described, resuspended in 0.2% PBST and filtered through a 20µm strainer (pluriSelect 431002040) for flow cytometry.

Antibodies used: p65^Ab^ (BioLegend Poly6226, cat #622601), p65^inCITE Ab^ (BioLegend Poly6226, #622601), c-Fos^inCITE Ab^ (BioLegend Poly6414, cat# 641401), NeuN^inCITE Ab^ (BioLegend 1B7, cat #834502), PU.1^inCITE Ab^ (BioLegend 7C2C34, cat #681307).

### inCITE antibodies

Pure clones of all antibodies were conjugated with the TotalSeq^TM^-A format (BioLegend).

### InCITE-seq

Antibody stained nuclei were resuspended in PBS with 3mM MgCl_2_, then filtered through a 10µm filter. Nuclei were counted in a hemocytometer chamber by eye, and promptly loaded onto a Chromium single-cell V3 3’ chip (10X Genomics) according to the manufacturer’s protocol for GEM formation. For the HeLa experiment, a single V3 3’ 10x channel was loaded with 10,000 nuclei of NT and 10,000 nuclei of TNF *α* treated sample with nucleus hashing. For the mouse hippocampus experiment, two V3 3’ 10x channels were loaded, each channel with 30,000 nuclei of a 1:1 mix of nucleus-hashed PBS sample and a kainic acid sample, such that in total, *n=*2 PBS sample and *n*=2 kainic acid treated sample were loaded.

After GEM formation, simultaneous reverse crosslinking and reverse transcription were conducted by incubating GEMs at 53°C for 45 min followed by 85°C for 5 min. Samples were stored at - 20°C until subsequent GEM recovery and cDNA amplification. cDNA was amplified using the standard 10X Genomics single cell 3’ V3 protocol (10X Genomics) with both HTO and ADT PCR additive primers included in the AMP mix at 0.1µM and 0.2µM, respectively. After amplification, antibody-oligo derived cDNA fragments were separated from mRNA derived cDNA through SPRI-based size selection by incubating cDNA amplification product in 0.6x SPRIselect (Beckman Coulter B23319) for 5 min at room temperature. As a result, antibody-oligo derived cDNA is contained in the supernatant, while mRNA derived cDNA remains on SPRIselect beads. Supernatant containing antibody-oligo derived cDNA was removed and separately stored for HTO/ADT library construction. SPRIselect containing mRNA derived cDNA (WTA) were washed two times in 80% ethanol, eluted into 40µL of Elution Buffer and stored for gene expression library construction.

For gene expression library construction, 10µL of cleaned WTA was used to construct final sequencing libraries using manufacturer specific enzymatic fragmentation, adaptor ligation, and sample index attachment. and the final sequencing library was eluted in 30µL of Elution Buffer according to the standard 10X Genomics single cell 3’ V3 protocol. Samples were stored at -20°C until library quantification and sequencing.

For HTO/ADT library construction, cDNA amplification supernatant containing antibody-oligo derived cDNA was mixed in 1.4x SPRIselect and incubated at room temperature for 5 min. Mixture was cleared on a magnet for 5 min before two standard 80% ethanol washes were performed and the sample was eluted into 24µL of ddH_2_O. Afterwards, 6µL of eluted solution was added to one of two PCR solutions containing either unique HTO or ADT index primer mix, and NEBNext 2x Mastermix (New England BioLabs M0541L) in order to construct separate HTO and ADT libraries. Library construction was completed through a PCR amplification protocol with the following conditions: 98°C for 5 min, 21 cycles at 98°C for 2 sec and 72°C for 15 sec, and a final 72°C for 1 min. PCR products were then purified through 2.0x SPRI 10 min incubation and 5 min magnetic separation incubation, and two 80% ethanol washes. Purified products were eluted into 20µL of EB. Samples were stored at 4°C until library quantification and sequencing.

Gene expression, HTO and ADT libraries were quantified using standard Qubit (ThermoFisher Q32853) and Agilent TapeStation (Agilent G2991AA) methods to check for library size. Optimal library size was ∼420bp for gene expression and ∼180bp for ADT/HTO. Libraries were pooled and sequenced on the NextSeq500 platform (Illumnia) (Read 1: 28 cycles, Index 1: 8 cycles, Read 2: 55 cycles).

### InCITE-seq data pre-processing

Sequencing data were processed with Cumulus version 1.0 using cellranger v4.0.0^99^. Reads from demultiplexed FASTQ files were aligned to pre-mRNA annotated genomes of the mouse mm10 and human GRCh38 reference genome, similarly to previously described^100^. Hashed nuclei were demultiplexed using DemuxEM^46^ with min_num_genes=10, min_num_umis=1, min_signal_hashtag=1; Nuclei with ambiguous treatment assignment from demultiplexing with demuxEM (*i.e.*, nuclei not assigned as NT or TNFα among HeLa nuclei, or PBS or KA among mouse hippocampus nuclei) were discarded (5.9% and 2.7% for HeLa and mouse, respectively). For mouse hippocampus data, raw counts across both channels were concatenated. Gene expression count matrices were subsequently analyzed by Scanpy (v1.6.0)^101^. Unspliced pre-mRNA and spliced mRNA counts were generated from BAM alignment files with Velocyto v0.17.17^17^ and added as layers to the anndata object. Protein ADT counts corresponding to each cell barcode were also merged.

### Analysis of gene expression

For HeLa cell data, gene expression matrices were filtered to retain genes found in at least 10 nuclei, and nuclei with at least 500 genes and at most 5,000 UMI counts, resulting in a matrix of 10,014 nuclei with 13,942 genes across conditions. For mouse hippocampus data we retained genes found in at least 3 nuclei, and nuclei with at least 30 genes and at most 900 or 1,200 UMI counts for batch 1 or 2, respectively; additionally, nuclei with mitochondrial gene content >5% and hashtag oligo counts (HTO counts) greater than 5,000 were removed, resulting in a matrix of 21,583 nuclei with 16,484 genes. Gene counts were normalized within each nucleus, then log normalized as ln(X+1).

### Normalizing protein expression

Protein abundances as measured by antibody-derived tag (ADT) were first normalized by nucleus hashtag oligo counts (HTO), with the addition of a pseudo-count: nADT = (ADT + 1)/HTO. These normalized nADT counts were then scaled to centered log ratios, resulting in nuclear Centered Log Ratios (“nCLR”): nCLR = nADT/(Π_i_ nADT_i_)^1/n^, where the denominator is the geometric mean.

### Clustering mouse hippocampus snRNA-seq data

4,954 variable genes on log normalized counts were selected using Scanpy’s highly_variable_genes function (min_mean=0.001, max_min=0.15, min_disp=0.25), log counts were scaled, and UMI counts and mitochondrial content were regressed out using Scanpy’s regress_out function. Dimensionality reduction was performed using the variable genes via PCA in Scanpy, followed by a python implementation of Harmony^102^ to correct for batch based on 10x channels and to regress out differences due to treatment (PBS and KA). Nearest-neighbor graph was constructed with *k=*10 neighbors and top 30 principal components, clustered with the Leiden algorithm in Scanpy, and embedded using UMAP^103^ in Scanpy.

### Genes associated with p65 protein levels in HeLa

Gene expression was modeled using a generalized linear model with a negative binomial fit, as follows: *Y_i_ ∼ p65 + G2M_score + S_score + log(UMI) + log(HTO)*, where *Y_i_* is the log normalized, unscaled ln(X+1) counts for gene *i, p65* is the p65 protein level in units of nCLR, G2M_score and S_score are cell cycle scores calculated with score_genes_cell_cycle in Scanpy using previously defined genes^104^, and *log(UMI)* and *log(HTO)* are natural log counts of unique RNA molecular identifiers and nucleus hashtag oligos. Significance was established at a false discovery rate (FDR) of 1% after Benjamini-Hocberg correction using the statsmodels package in python.

### Gene Ontology analysis

Gene sets were queried using scanpy’s queries.enrich module, a wrapper around gprofiler^105^, to identify Gene Ontology (GO) Biological Processes. P-values were calculated using a hypergeometric test.

### Genes globally associated with protein levels in mouse hippocampus

We implemented a two-step mixed linear model using the statsmodel package in Python. First, we modeled gene expression as the following, to regress out effects of treatment and cell type: *Y_i_ ∼ C(cluster) + C(treatment) + C(cluster)*C(treatment) + log(UMI) + log(HTO) + (1|B),* where *Y_i_* is the scaled z-score of the ln(X+1) counts for gene *i* across all nuclei*, C(cluster)* is a categorical variable indicating cluster membership (cell type), *C(treatment)* is a categorical variable indicating PBS or KA treatment, *log(UMI)* and *log(HTO)* are natural log counts of unique RNA molecular identifiers and nucleus hashtag oligos, and *B* is a categorical variable denoting the 10x channel batch. We then modeled the residuals of each gene, *rY_i_*, as a linear combination of the four proteins, accounting for batch: *rY_i_ ∼ NeuN + c-Fos + p65 + PU.1 + (1|B)*. Genes found in at least 20 nuclei were used for analysis, and significance was established at a false discovery rate (FDR) of 5% after Benjamini-Hochberg correction. For identifying treatment-dependent global effects, we implemented the same two-step strategy, but using nuclei populations that were only in PBS treated or KA treated.

### Genes associated with protein levels in mouse hippocampus within excitatory neurons and astroglia

Similar to the global approach, we first regressed out treatment within each cluster: *Y_i,c_ ∼ C(treatment) + log(UMI) + log(HTO) + (1|B)*, where all variates are the same as previously described, except that Y_i,c_ is the scaled z-score of the ln(X+1) counts for gene *i* in cluster *c*. We then modeled the residuals of each gene as a linear combination of the four proteins, accounting for batch: *rY_i_ ∼ NeuN + c-Fos + p65 + PU.1 + (1|B)*. Genes found in ≥10 nuclei were used for analysis, and significance was established at a false discovery rate (FDR) of 5% with Benjamini-Hochberg correction.

### Identifying cell type-specific gene programs with non-negative matrix factorization

We identified gene programs from the RNA profiles of excitatory neurons or astroglia, using a subset of genes for each cell type (a combination of highly variable genes identified for clustering and c-Fos and p65-associated genes within each cell type, with manual removal of the highly expressed and variable lncRNA Gm42418), via non-negative matrix factorization using the sklearn package in Python (NMF function; random seed, random_state=0, L1 regularization with l1_ratio=1, alpha=0). Four and three programs were identified for EX neurons and astroglia, respectively.

### Cell type-specific genes associated with protein levels in mouse hippocampus after KA treatment

We implemented the following mixed linear model for each cell cluster *c* using the statsmodels package in Python: *Y_i,c_ ∼ NeuN + c-Fos + p65 + PU.1 + log(UMI) + log(HTO) + (1|B)*, where: *Y_i.c_* is the scaled z-score of the ln(X+1) counts for gene *i* in cluster *c, NeuN, c-Fos, p65, PU.1* are protein levels in units of nCLR, *log(UMI)* and *log(HTO)* are natural log counts of unique RNA molecular identifiers and nucleus hashtag oligos, and *B* is a categorical variable denoting the 10x channel batch. Protein levels (nCLR) were first scaled using the Python package sklearn preprocessing. Genes found in ≥10 nuclei of that cluster were used for analysis, and significance was established at a false discovery rate (FDR) of 5% with Benjamini-Hochberg correction.

### Transcription factor motif enrichment in DEGs

We used differentially accessible regions (DARs) of the mouse hippocampus profiled in saline treated and kainic acid treated (1hr post treatment) previously defined by Fernandez-Albert et al^75^. Nearest DARs located more than 1kb from the transcriptional start site (upstream) or transcriptional termination site (downstream) of DEGs associated with inCITE-seq measurements of c-Fos or p65 were considered as enhancers and were used for motif enrichment analysis. To find enriched motifs, we scanned a given set of differentially accessible peaks for all DNA-binding motifs in the cis-bs (http://cisbp.ccbr.utoronto.ca) and JASPAR2018_CORE_vertebrates_non-redundant (http://jaspar2018.genereg.net) databases. We then tested for the probability of our observed motif frequency using a hypergeometric test, with a null model based on random sampling of all ATAC-seq peaks matching for GC content. Motifs with *P*<10^-5^ were considered significant.

### Code availability

Code available on https://github.com/klarman-cell-observatory/inCITE-seq.

## Acknowledgments

We thank L. Gaffney for assistance with figure preparation, P. Thakore for coining the acronym inCITE-seq, A. Rubin for critical feedback on the manuscript, E. Fiskin, S. Simmons, O. Ursu, C. Smillie, D. Silverbush, E. Torlai-Triglia, and members of the Regev lab for helpful discussions, and the Broad Institute Flow Cytometry Core facility. This research was supported by NIH/NHGRI CEGS grant 5RM1 HG006193. A.R. was a Howard Hughes Medical Institute Investigator (until July 31, 2020).

## Author Contributions

H.C. conceived and designed the study with guidance from A.R. C.N.P. designed and performed mouse experiments with guidance from D.A. H.C. and E.M. developed and performed inCITE-seq experiments, with buffer optimization input from F.C. and early stage experimental support from J.W. C.N.P. and E.M. conducted immunohistochemistry. D.P. conducted 10x experiments and constructed sequencing libraries. B.Y. conjugated inCITE antibodies. H.C. analyzed and interpreted the data, with help from E.H. on cis regulatory motif enrichment analysis, and supervision from A.R. A.R. provided project oversight and funding. H.C. and A.R. wrote the manuscript with input from all authors.

---

**Supplementary Fig. 1.**
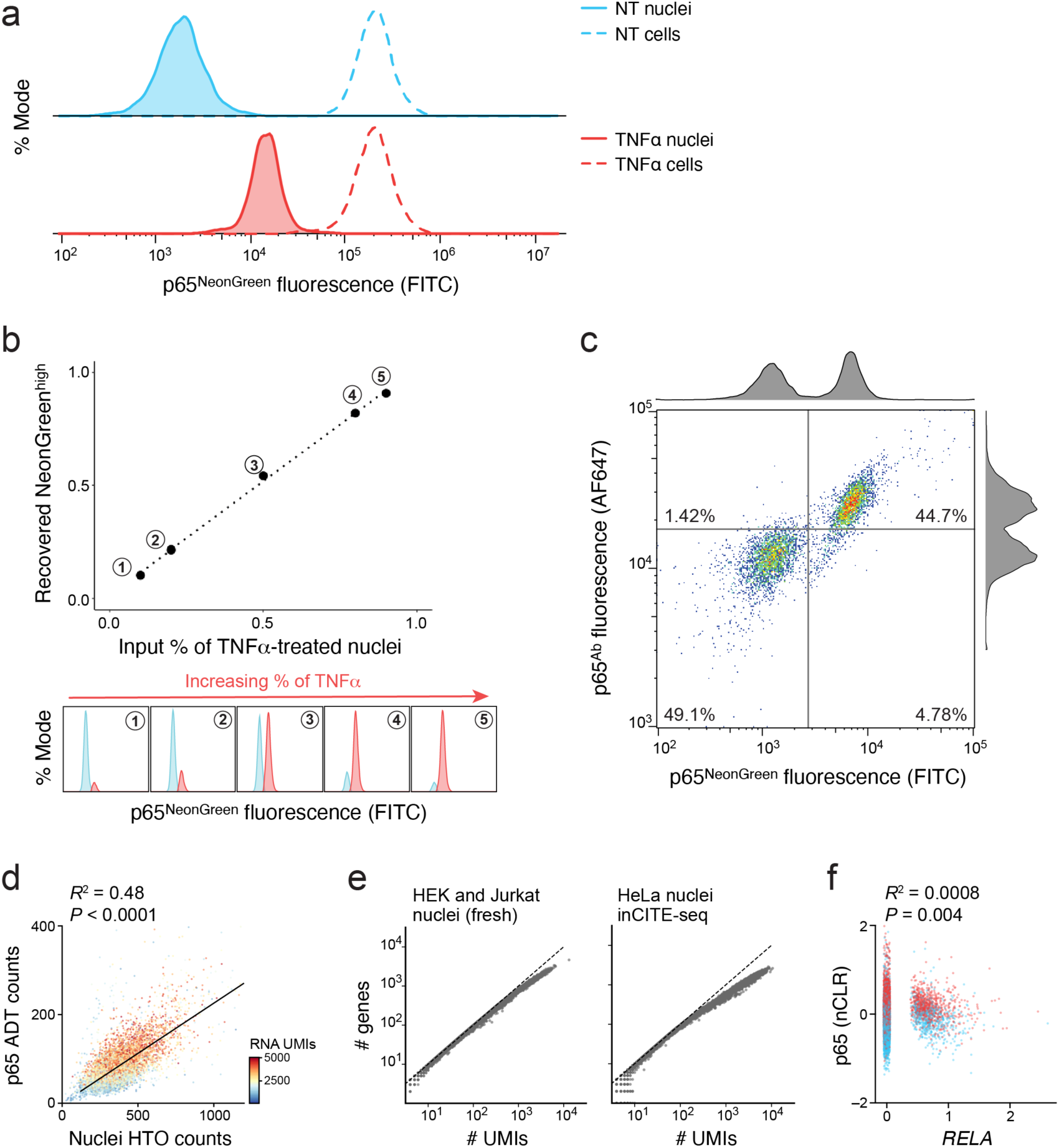
Optimization of intranuclear antibody staining in HeLa cells. **a.** p65 nuclear localization following TNFα treatment. Distribution of p65-mNeonGreen reporter fluorescence (x axis; % mode of singlet nuclei, y axis) measured by flow cytometry of nuclei (solid line) and cells (dashed line) that were untreated (“NT”, blue) or collected 40 min after TNFα treatment (red). **b.** Flow cytometry distinguishes low and high nuclear p65 signals across a range of NT and TNFα nuclei ratios. Top: Measured proportion of NeonGreen^high^ to mNeonGreen^low^ nuclei by flow cytometry (x axis) and the actual proportion of input nuclei stimulated with TNFα to untreated nuclei (TNFα:NT ratio, x axis). Bottom: Corresponding flow cytometry distributions of mNeonGreen^high^ (red) and mNeonGreen^low^ (blue). **c.** Agreement between antibody- and fluorescence-based estimated of p65. Antibody-derived (*y* axis, p65 antibody signal amplified by Alexa Fluor 647 conjugated secondary) and mNeonGreen (x axis) measurements of p65 in an equal mixture of nuclei from untreated (NT) and TNFα stimulated nuclei. Histograms: marginal distributions. **d.** Relation between nuclei hashtag oligonucleotide counts and p65 antibody-derived tag counts. HTO (x axis) and p65 antibody-derived tag counts (ADT; y axis) for 10,014 nuclei from NT and TNFα HeLa populations, colored by number of RNA UMIs. Top left: Pearson R^2^ and associated P-value. To control for this relation, we subsequently normalize protein ADT counts by nuclei HTO counts for nucleus bead barcode (**Methods**). **e.** InCITE-seq RNA library complexity compares well to snRNA-seq. Number of detected genes (y axis) and transcripts (UMIs) (x axis) (without filtering) in snRNA-seq from HEK and Jurkat cell lines (McGinnis et al. 2019) (left) and from inCITE-seq of Hela cells (right). **f.** Low correlation between p65 protein and RNA levels. p65 protein levels (*y* axis, nCLR, **Methods**) and RNA levels (*RELA*, x axis, log counts) from InCITE-Seq. Top left: Pearson R^2^ and associated P-value.

**Supplementary Fig. 2.**
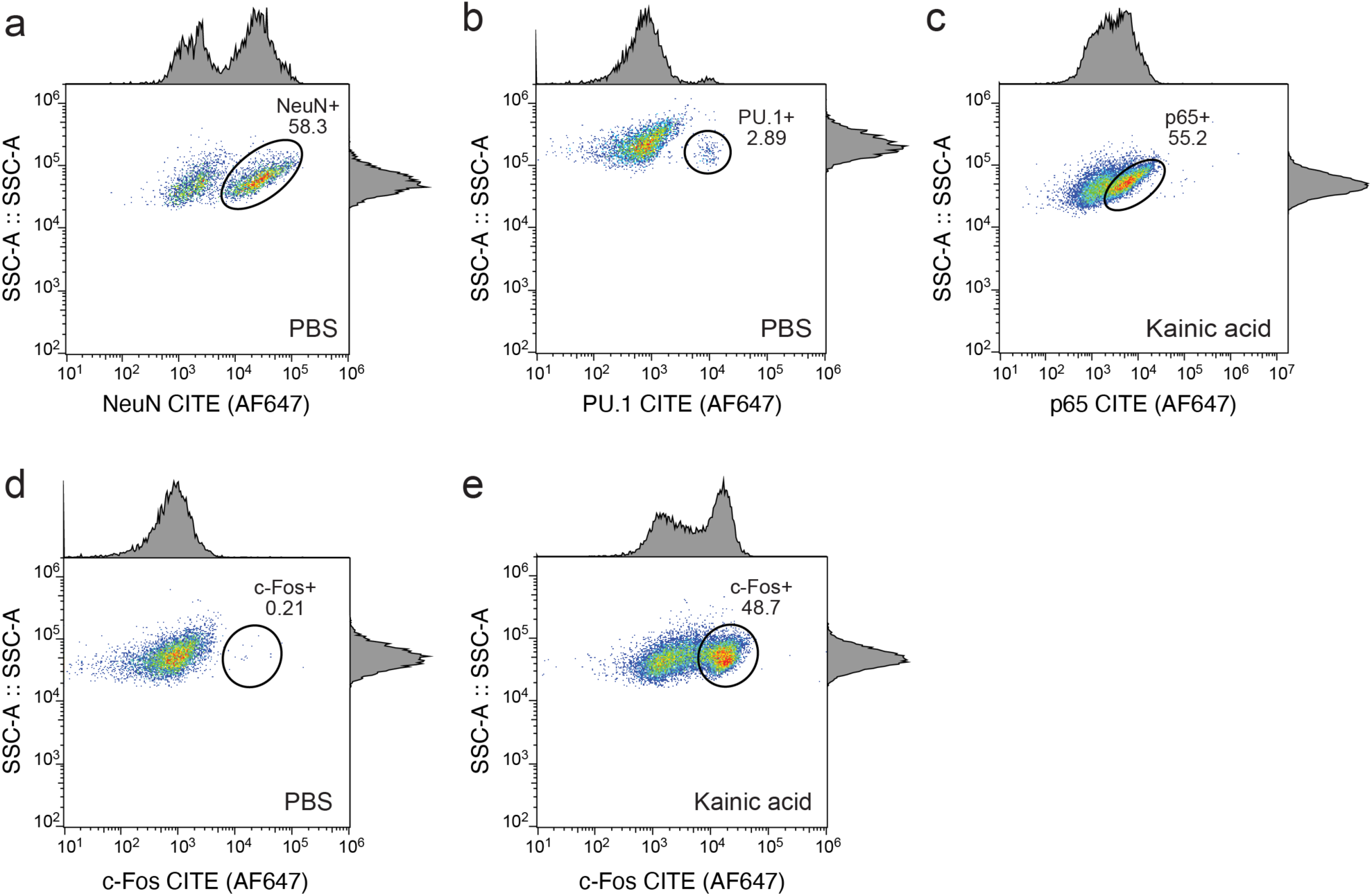
Flow cytometry of inCITE targets on nuclei from the mouse hippocampus. Flow cytometry of nuclei populations from the mouse hippocampus after primary antibody stains with inCITE antibodies targeting NeuN in PBS (**a**), PU.1 in PBS (**b**), p65 in kainic acid (KA) (**c**), and c-Fos in PBS (**d**) and kainic acid (KA) (**e**) treated mice, followed by Alexa Fluor 647- conjugated secondary stain (*x* axis), compared to side scatter (*y* axis). Histograms show marginal distributions. Ovals mark NeuN^high^ (A, 58.3%), PU.1^high^ (B, <3%), p65^high^ (55.2%), c-Fos^high^ (0.21% in PBS (**d**), and 48.7% after KA treatment (**e**)).

**Supplementary Fig. 3.**
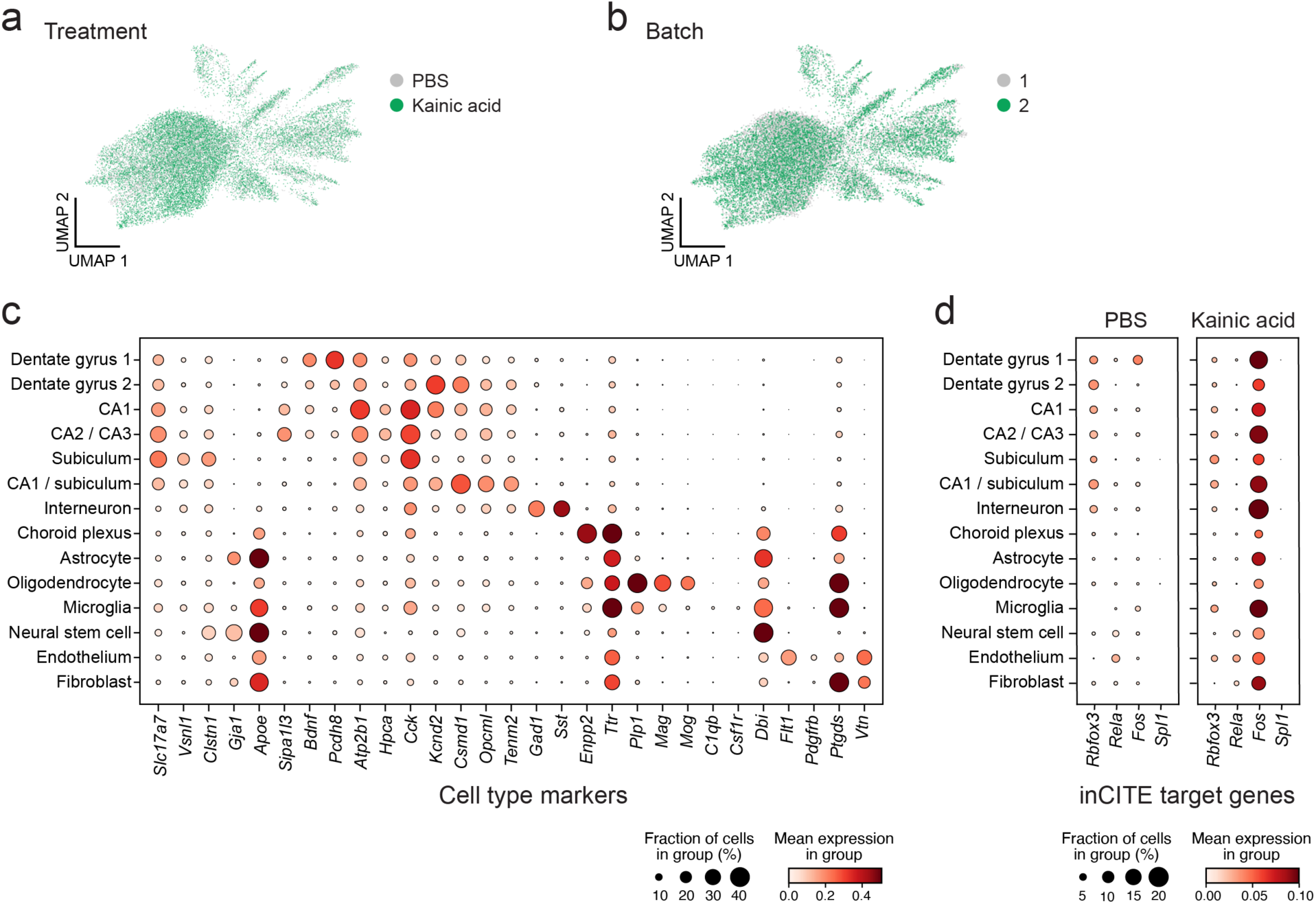
Gene expression patterns in mouse hippocampus nuclei. **a,b.** Controlling for treatment and batch. UMAP embedding (as in Fig. 2b) of single nucleus RNA profiles from InCITE-seq of the mouse hippocampus, after regressing treatment and batch (**Methods**), colored by treatment (PBS, gray; kainic acid, green), and batch (batch 1, gray; batch 2, green; each batch is one channel loaded with a multiplexed pool of a PBS and a KA treated sample with nucleus hashing; **Methods**). **c.** Cell type marker genes. Mean expression of log normalized counts (dot color) and proportion of expressing cells (dot size) of marker genes (columns) used for annotating cell subset clusters (rows). **d.** RNA expression of genes encoding inCITE protein targets. Mean expression of log normalized counts (dot color) and proportion of expressing cells (dot size) for RNAs encoding the inCITE-seq protein targets (*x* axis) in each cluster (*y* axis) in PBS (left) and KA (right) treatment. *Rbfox3* encodes NeuN, *Spi1* encodes PU.1, *Fos* encodes c-Fos, and *Rela* encodes p65.

**Supplementary Fig. 4.**
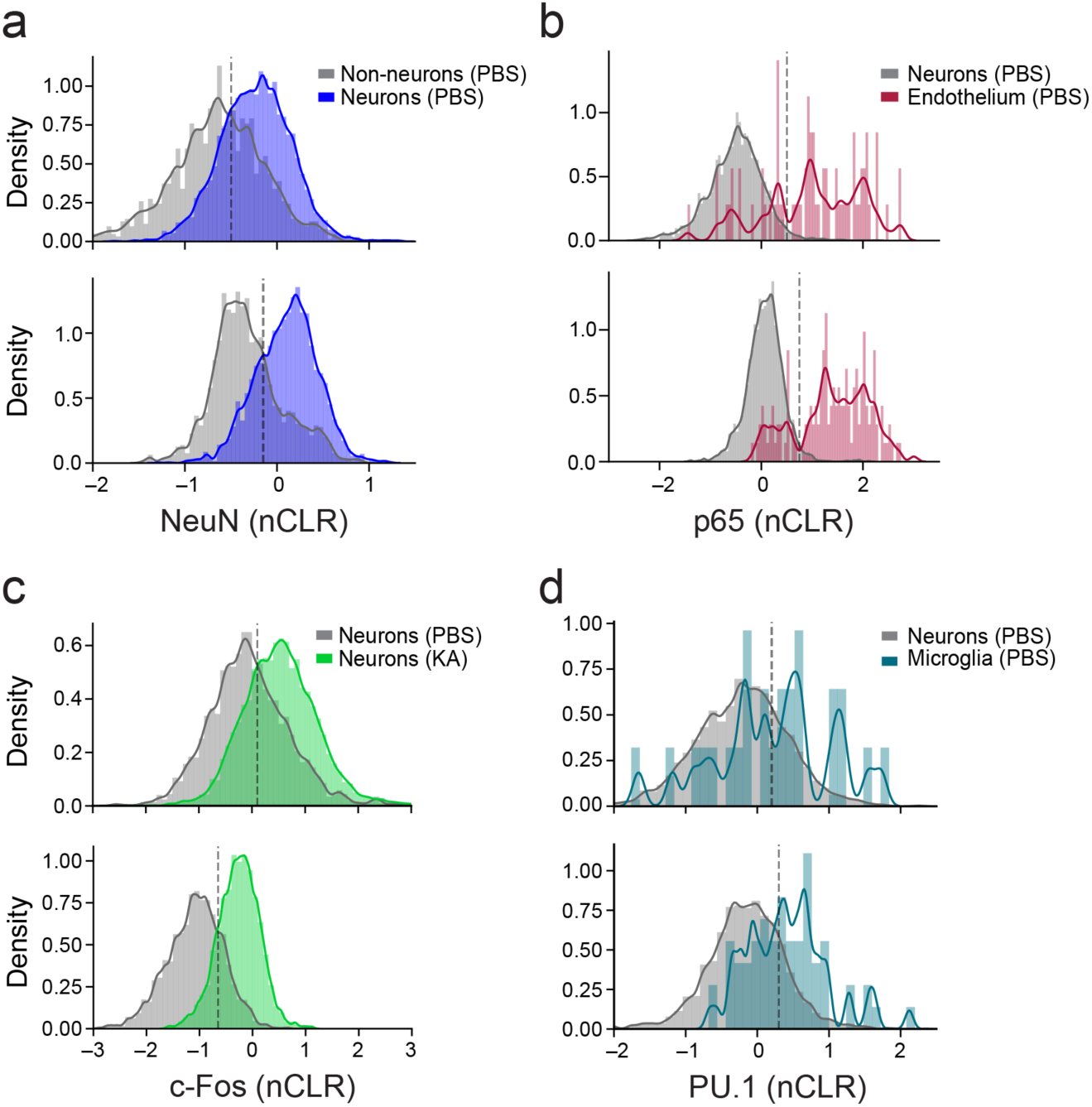
Batch specific inCITE-seq measures of protein levels. Distribution (kernel density estimates, curve) of protein levels (*x* axis, nCLR) of NeuN (**a**), p65 (**b**), c-Fos (**c**), or PU.1 (**d**) in each batch (top: batch 1; bottom: batch 2) in a subset of interest (color) and appropriate background set of nuclei (grey). Dashed line: Batch-specific threshold used to partition protein level as high vs. low.

**Supplementary Fig. 5.**
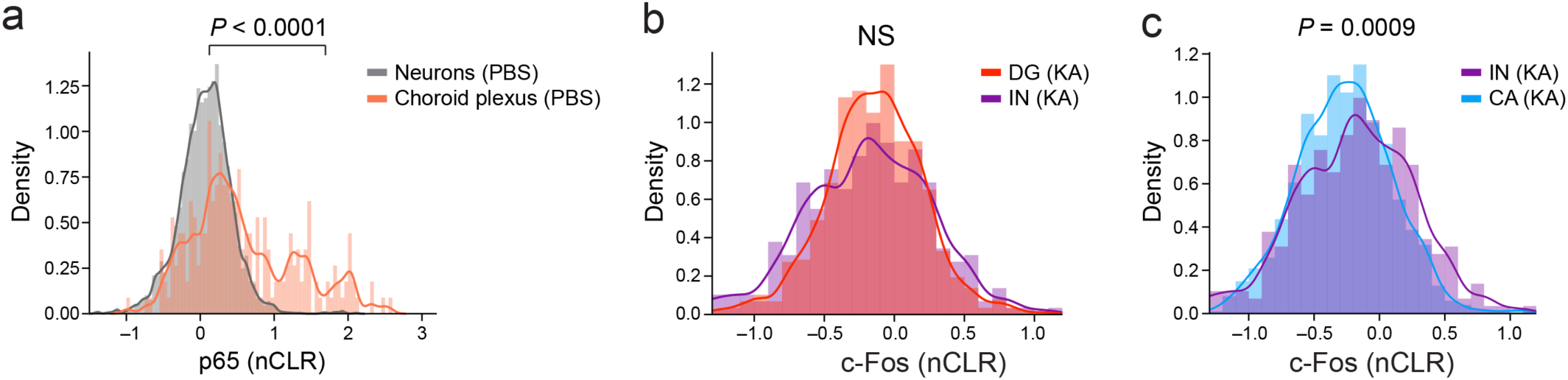
Differences in protein levels between cell types measured by inCITE-seq. Distribution (kernel density estimates, curve) of protein levels (nCLR) of p65 (**a**) and c-Fos (**b,c**) from single batches. **a.** p65 levels are higher in subpopulations of the choroid plexus (orange) compared to neurons (grey) in PBS treated mice. **b.** c-Fos levels are indistinguishable in KA treated mice between dentate gyrus neuron clusters (DG1 and DG2, red) and interneurons (purple). **C.** c-Fos levels are higher in KA treated mice in interneurons (IN, purple) than in CA neuron clusters (CA1 and CA2/3). *P*-values, t-test. NS – not significant.

**Supplementary Fig. 6.**
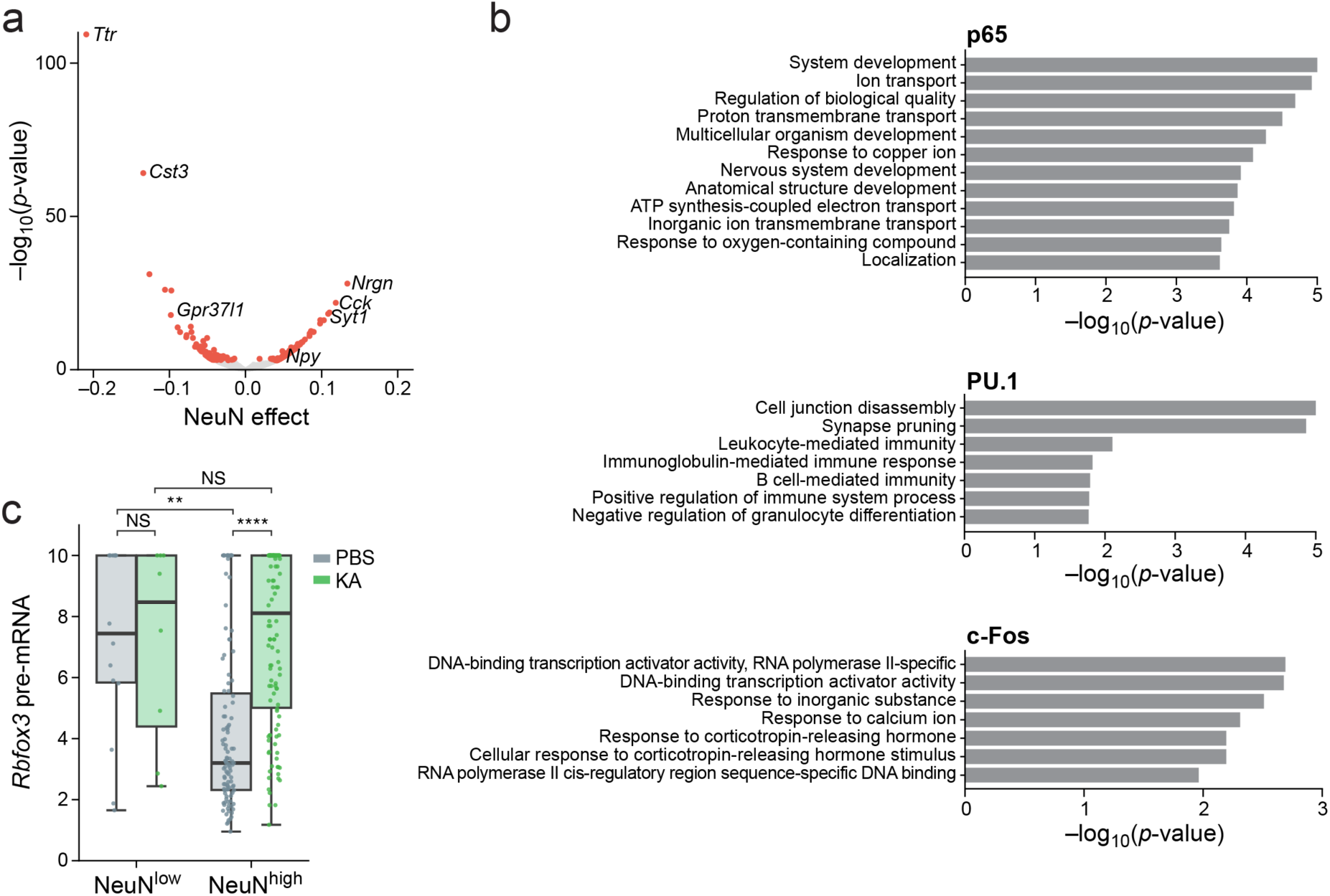
Protein effects on global gene expression. **a.** Genes associated with NeuN. Effect size (x axis) and associated significance (y axis, -log_10_(P-value)) for the association of each gene (dots) with NeuN by a model of gene expression as a linear combination of the four inCITE-seq target proteins after regressing out treatment and cell type (**Methods**). Select genes are labeled. Colored dots: Benjamini-Hochberg FDR <5%. **b.** Functional gene sets enriched with TF associated genes. Enrichment (-log_10_(P-value), *x* axis, hypergeometric test) of Gene Ontology (GO) terms (*y* axis) in genes significantly associated (from top to bottom) with p65 (131 genes), PU.1 (16 genes), and c-Fos (16 genes). **c.** Relation between unspliced, pre-mRNA and nuclear protein levels of NeuN. Distribution of pre-mRNA levels (Z score of log-normalized counts, y axis) in nuclei with high or low levels (**Supplementary Fig. 4**) of NeuN (x axis) after PBS (gray) or KA (green) treatment. Boxplot shows the median, with ends representing 25% and 75% quartiles and whiskers at 1.5 interquartile range below 25% or above 75% quartiles. Dots correspond to individual nuclei with non-zero pre-mRNA levels. **P<10^-2^, **** *P*<10^-4^, t-test. NS – not significant.

**Supplementary Fig. 7.**
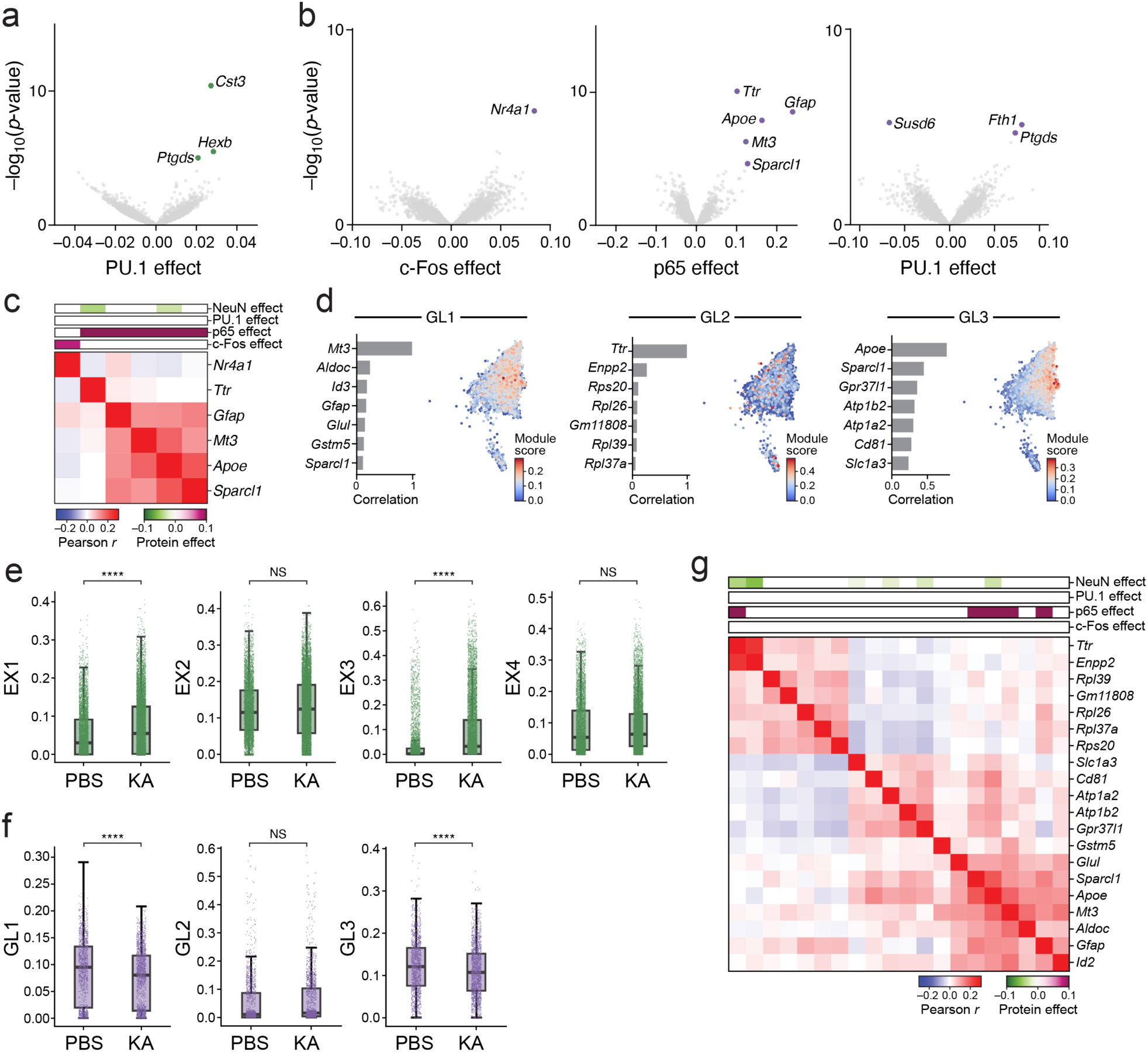
Genes and modules associated with TFs within excitatory (EX) neurons and astroglia (astrocyte and microglia). **a,b.** Genes associated with each TF within excitatory neurons or astroglia. Effect size (x axis) and significance (y axis, -log_10_(P-value)) for genes (dots) globally associated with each TF (label at bottom) by a model of gene expression as a linear combination of proteins after regressing out treatment (**Methods**) applied in all excitatory neurons (**a**) or astrocytes (**b**). Select genes are labeled. Colored dots: Benjamini-Hochberg FDR<5%. **c.** Key genes associated with p65 and c-Fos in astroglia. Pearson correlation coefficient (red/blue colorbar) between the expression profiles of genes (rows and columns) across astroglia cells, for genes significantly (FDR<5%) associated positively (purple) or negatively (green) with either c-Fos or p65 in astroglia, ordered by hierarchical clustering. Top bars: Effect size of each TF from the linear model in astroglia (**Methods**). **d.** NMF-derived gene programs within astroglia. Right: UMAP embedding of astroglia nuclei RNA profiles from inCITE-seq, colored by NMF module score (colorbar). Left: Top 7 module genes (y axis) and their Pearson correlation coefficient with module scores (x axis). **e,f.** Treatment effect on gene programs. Program score (y axis) for nuclei (dots) from PBS or KA treated mice (x axis) for excitatory neuron programs (**e**) and astroglia programs (**f**). \*\*\*\**P*<10^-4^, t-test. NS – not significant. **g.** Pearson pairwise correlation coefficient (red/blue) between gene expression profiles of each of the top 7 genes of the astroglia NMF programs (rows), ordered by hierarchical clustering. Top bars (purple/green): significant effect size of each protein from the linear model.

**Supplementary Fig. 8.**
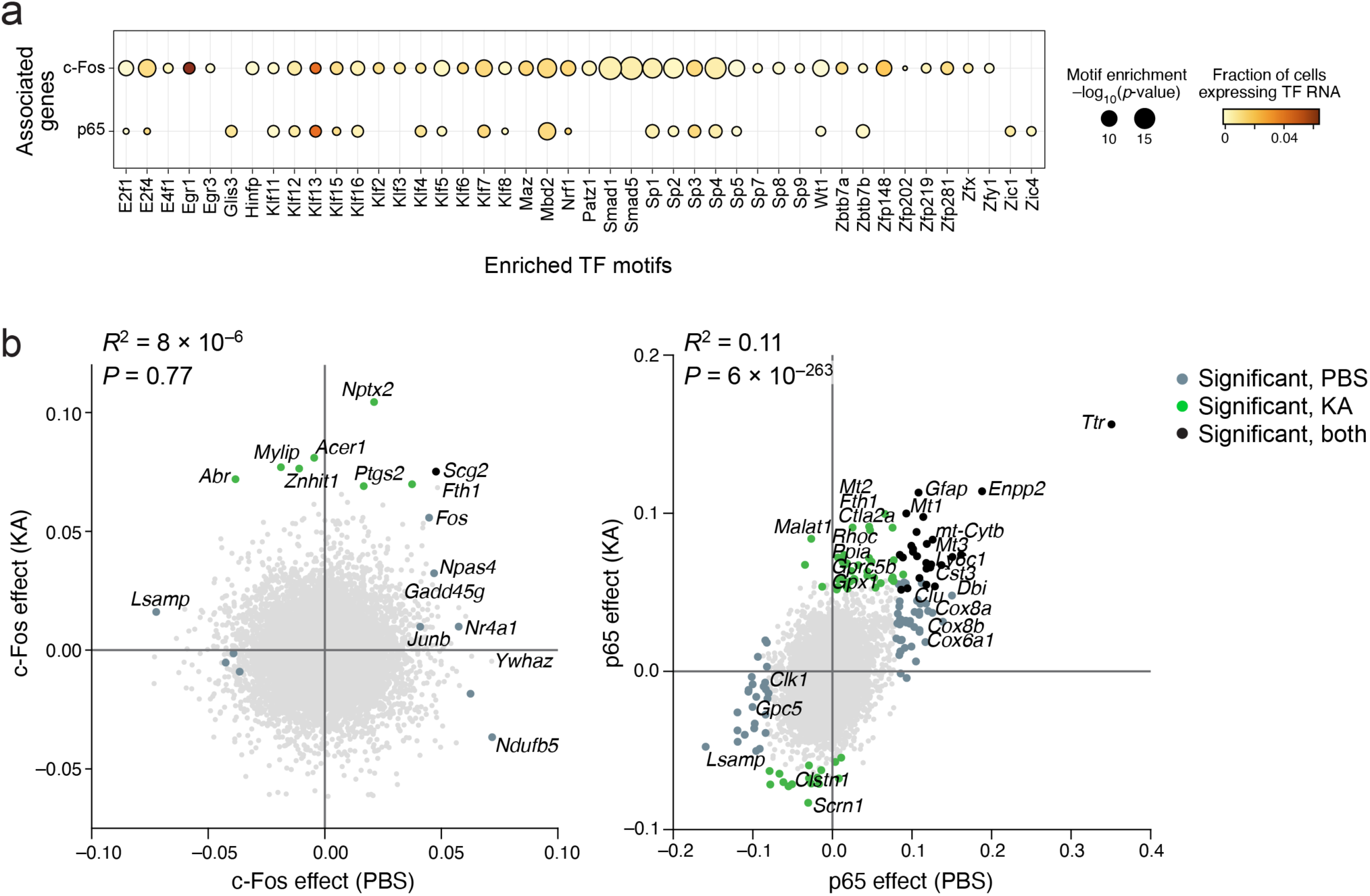
Treatment-dependent *cis*-regulatory elements and TF-associated genes. **a.** TF motifs enriched in enhancers of c-Fos and p65 associated genes. Significance (dot size, - log_10_(P-value) of enrichment, hypergeometric test) for TFs (columns) whose motifs are significantly enriched in enhancers of genes associated with c-Fos or p65 (rows) in excitatory neurons compared to enhancers of non-associated genes in KA treated samples, and proportion of excitatory neuron nuclei expressing the cognate TF’s RNA (dot color). **b.** Treatment-dependence of gene association with c-Fos and p65. Global effect size of genes (dots) associated with c-Fos (left) and p65 (right) after PBS (x axis) or KA treatment (y axis), by a model of gene expression as a linear combination of the four proteins after regressing out treatment and cell type (**Methods**). Colored dots: genes with significant (Benjamini-Hochberg FDR<5%) coefficients in PBS (grey), KA (green), or both (black). Select genes are labeled. Top left: R^2^ and associated *P* value.

